# Risk modelling of single cell populations revealed the heterogeneity of immune infiltration in hepatocellular carcinoma

**DOI:** 10.1101/2021.06.26.450025

**Authors:** Lu Wang, Yifan Chen, Fengbiao Mao, Zhongsheng Sun, Xiangdong Liu

## Abstract

Hepatocellular carcinoma (HCC) is one of the most common primary hepatic malignancies. E2F transcription factors play an important role in the tumorigenesis and progression of HCC, mainly through the RB/E2F pathway. Prognostic models for HCC based on gene signatures have been developed rapidly in recent years; however, their discriminating ability at the single cell level remains elusive, which could reflect the underlying mechanisms driving the sample bifurcation. In this study, we constructed and validated a predictive model based on E2F expression, successfully stratifying HCC patients into two groups with different survival risks. Then we used a single-cell dataset to test the discriminating ability of the predictive model on infiltrating T cells, demonstrating remarkable cellular heterogeneity as well as altered cell fates. We identified distinct cell subpopulations with diverse molecular characteristics. We also found that the distribution of cell subpopulations varied considerably across onset stages among patients, providing fundamental basis for patient-oriented precision evaluation. Moreover, single-sample gene set enrichment analysis revealed that subsets of CD8^+^ T cells with significantly different cell adhesion level could be associated with different patterns of tumor cell dissemination. Therefore, our findings linked the conventional prognostic gene signature to the immune microenvironment and cellular heterogeneity at the single-cell level, thereby providing deeper insights into the understanding of HCC tumorigenesis.

## Introduction

Hepatocellular carcinoma (HCC), the most prevalent type of primary liver cancer, accounts for up to 90% of all liver cancers (1). It typically develops on a background of chronic liver inflammation or cirrhosis, with hepatitis B viral infections being the primary risk factor for its development (2). Although great efforts have been made in the early diagnosis and treatment methods of HCC, the prognosis of HCC patients still requires much improvement. In addition, the current TNM staging system and other traditional clinical models may be useful for prognostic assessment, but are still not ideally accurate in the grading of patients for surgical treatments. Over the past decade, immunotherapeutic approaches have changed the landscape of clinical decision-making for HCC patients, albeit with rather suboptimal response rates (3, 4). The high heterogeneity at the cellular and molecular levels may pose a considerable obstacle to improving the therapeutic efficacy of HCC. Although diverse predictive models for prognosis are capable of classifying patients into different risk groups, which corroborates inter-patient heterogeneity, the factors underlying the discriminative effects of these models have not been studied. Thus, exploring the tumor microenvironment and dissecting the developmental dynamics of immune cells are pivotal for understanding the heterogeneous pattern of antitumor immune responses in HCC patients. Single-cell sequencing methods have emerged as cutting-edge techniques for analyzing the complex tumor ecosystem and characterizing immune microenvironmental patterns, and several previous studies have demonstrated the feasibility of investigating the tumor immune phenotypes at a single-cell resolution for HCC (5–7).

Unlike cells in other organs or tissues that are regenerated by stem cells, hepatocytes are characterized by an impressive ability to maintain their remarkable proliferative potential, which compensates for cell loss or is in response to hyperplasia (8). Members of the E2F transcription factor family are important in the regulation of hepatocyte proliferation, differentiation, apoptosis, and tumorigenesis in collaboration with cyclin-dependent kinases (CDKs) and the RB family of proteins (9–13). As previously demonstrated, dysfunction of the CDK-RB-E2F axis would leads to an aberrant cell cycle and the inhibition of apoptosis, and thus initiating pathological processes of HCC (10, 14, 15). Aberrant expression of E2F family members are reported to be associated with poor survival and could be used to predict the prognosis of several tumor types (16–18).

Here, we aim to develop and validate a prognostic signature with E2F genes that play essential roles in the oncogenic processes of HCC. In addition, we intended to identify distinct subpopulations of T cells by applying the E2F signature on a singlecell transcriptome profile (5). We sought to delineate the differential cell composition, nonnegligible transcriptional heterogeneity, and inconsistent cell fate decisions between different subpopulations of infiltrating T cells. Additionally, we demonstrated a specific population of CD8^+^ T cells with prominently elevated cell adhesion, potentially resulting in a more effective inhibition of tumor dissemination than other cells. Our study offers novel insights into the underlying biology of varying clinical outcomes at the single-cell level and dissects intricate admixtures of immune cell populations, providing a new lens on the determinants of molecular and phenotypic heterogeneity in HCC biology.

## Materials and methods

### Differential gene expression analysis

Differential gene expression analyses were performed on the TCGA-LIHC and integrated GEO datasets. In the TCGA cohort, we included 369 HCC patients and 50 normal control samples. Four independent microarray studies (GSE76427, GSE136247, GSE107170, and GSE102079) were collected from the GEO database to generate two integrated profiles, including cohorts of HCC samples and normal controls. The batch effects from different studies were mitigated using the ComBat method implemented in the R sva package (19). The limma package was applied to calculate the fold changes of genes between HCC samples and normal controls, and genes with adjusted *P* value <0.05 and fold change >2 were identified as differentially expressed genes.

### Construction of the prognostic signature

The level III transcriptome profiles, including mRNA-seq data of 369 HCC patients and the corresponding clinical information from the TCGA-LIHC dataset were used to build and validate the prognostic signature. The clinical parameters of age, gender, family history of cancer, pathologic stage, histologic grade, Ishak fibrosis score, Child– Pugh grade, vascular invasion, alpha fetoprotein outcome, and residual tumor were evaluated in this study. After removing samples with incomplete survival information, 339 HCC patients were retained for subsequent analysis. The HCC patients were randomly assigned to a training set or a validation set at a ratio of 1:1 *(n* = 171 and *n* = 168 for training and validation sets).

To establish a clinically translatable prognostic gene signature, we initially performed univariate Cox proportional hazards regression analysis on the genes of the E2F family using expression data from the TCGA training set. Genes with an adjusted *P* value < 0.05 were considered significant and retained for subsequent stepwise multivariate Cox analysis. Finally, the prognostic-related gene signature was built, and the risk score derived from the gene signature was calculated as the sum of the expression level of each E2F gene multiplied by the corresponding regression coefficients obtained from the multivariate Cox analysis. The median risk score of the training cohort was chosen as the cutoff to stratify the patients into high- or low-risk groups for the TCGA HCC patients. Kaplan Meier survival analyses and the log-rank test were utilized to examine the predictive performance for this signature.

### External validation of the prognostic signature and construction of the quantitative nomogram

The GSE14520 dataset with 222 HCC patients and complete clinical information was retrieved from the GEO database. The risk score for each patient in the GSE14520 validation dataset was calculated based on the prognostic signature and stratified into high- and low-risk groups based on the median risk score derived from the TCGA training cohort. A Kaplan Meier survival analysis was carried out to assess the predictive ability.

To individualize the predicted results on the 1-year, 3-year and 5-year survival probability for HCC patients and further improve the predictive accuracy of our prognostic signature, we established a nomogram as a quantitative tool based on the risk score derived from the prognostic signature combined with common clinical characteristics. A Kaplan Meier survival analysis was performed and the C-index was taken as a measure of discrimination calculated to evaluate the capacity for the nomogram. Calibration plots were used to compare and display the agreement between the predicted results and observed outcomes with regard to the 1-year, 3-year and 5year overall survival for the HCC patients from the TCGA cohort.

### Estimation of the tumor-infiltrating immune cells in bulk-sequencing data

To demonstrate the differences in the proportions of diverse tumor-infiltrating immune cells between the high- and low-risk samples, we applied the CIBERSORT algorithm and LM22 signature matrix consisting of 547 genes to derive the immune infiltration status for 22 mature human hematopoietic populations in the HCC patients. The permutation test of 1,000 iterations was adopted to obtain the result of statistical significance followed by quantile normalization.

### Description of the single-cell RNA sequencing data

The single-cell transcriptome data of human infiltrating T cells in a gene expression matrix format was downloaded from the GEO database with the accession number GSE98638. T cells within this dataset were collected from peripheral blood, tumor tissues, and adjacent normal liver tissues and sequenced using the Illumina HiSeq platform. A total of 11 main clusters were described. C01_CD8 cluster: naive CD8^+^ T cells; C02_CD8 cluster: effector CD8^+^ T cells; C03_CD8 cluster: mucosal-associated invariant T cells (MAIT); C04_CD8 cluster: exhausted CD8^+^ T cells; C05_CD8 cluster: intermediate state CD8^+^ T cells between effector and exhausted CD8^+^ T cells; C06_CD4: naive CD4^+^ T cells; C07_CD4: peripheral T regulatory cells (Tregs); C08_CD4: tumor Tregs; C09_CD4: mixed state cells; C10_CD4: exhausted CD4^+^ T cells; C11_CD4: cytotoxic CD4^+^ T cells.

### Definition of Class I and Class II subpopulations

The coefficients from the previously constructed prognostic gene signature were applied on the centered single cell expression data to categorize the single cells. Unlike in bulk sequencing data, we did not consider the survival time and status of specific patients, instead we treated each single cell as a separate predicted instance. The median score of all single cells derived from the gene signature was used as the cutoff value. The cells with predicted scores above the cutoff were categorized as the Class I cells while the rest were classified as Class II cells.

### Pseudotime trajectory inference and branch-dependent gene expression analysis

The pseudotime developmental cell trajectory for the single-cell data was inferred by the Monocle 2 method (20). From the pseudotime analysis, MAIT cells were excluded owing to the distinct characteristics of the T cell receptor (TCR) (5). The DifferentialGeneTest function was used to identify the differentially expressed genes along pseudotime and only genes with an adjusted *P* value < 0.01 were retained to build the trajectory path. The identification of the cell fate decision points and branch-dependent gene expression analysis were carried out using the BEAM function implemented in Monocle 2.

## Results

### Essential role of the E2F regulators in HCC and the E2F predictive model of survival for HCC patients

To comprehensively identify essential regulatory factors that orchestrate the transcriptional architecture of HCC, we carried out differential gene expression analyses on the TCGA LIHC data set and integrated gene data set from four GEO expression profiles (Figs. 1A and 1B, Tables S1 and S2). Gene set enrichment analysis on both data sets revealed that the most enriched pathway for transcriptional activities in HCC was ascribed to E2F-target genes (Figs. 1C and 1D), indicating that the proper combination of the E2F transcription factors might be promising in constructing the prognostic model of HCC (21).

**Fig. 1.**
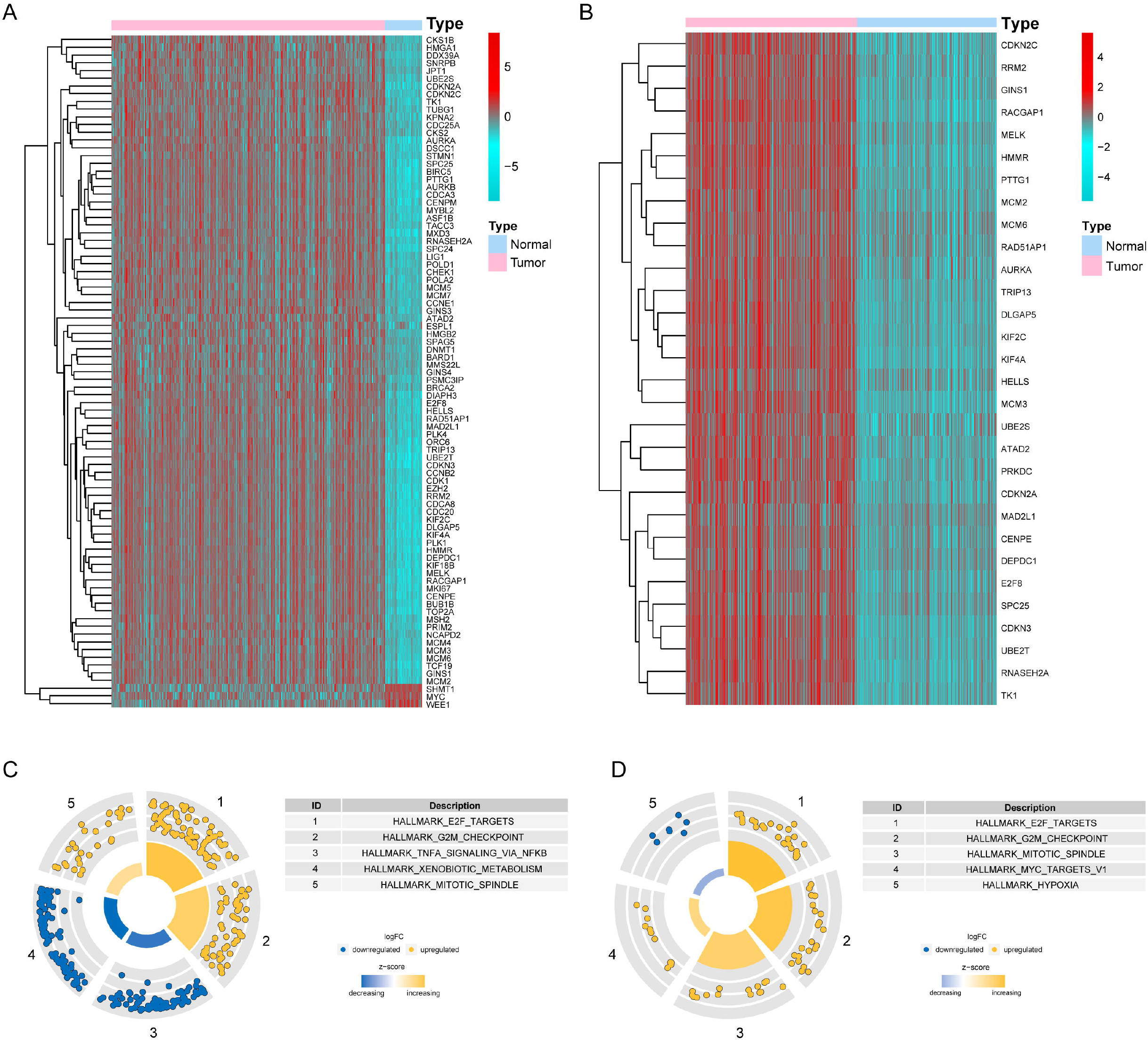
E2F transcription factors are essential regulators in HCC. **A-B**: Differential expression of E2F-target genes in the TCGA cohort (**A**) and the integrated GEO cohort (**B**). **C-D**: Gene set enrichment analysis showed the five most enriched biological processes for the differentially expressed genes in the TCGA cohort (**C**) and the combined GEO cohort (**D**).

Based on the expression data and clinical information from the TCGA LIHC cohort, we applied a Cox proportional hazards regression analysis on the E2F family members. After conducting a stepwise multivariate Cox analysis on the training cohort, E2F2 and E2F5 were identified to be significantly correlated with poor survival of the HCC patients (Table S3). Within the prognostic signature, a risk score was calculated for each patient according to the expression level and the corresponding regression coefficient of E2F2 and E2F5. Based on the median risk score for the training cohort, the HCC patients were categorized into high and low risk groups. We observed that patients in the low-risk group exhibited remarkably longer overall survival (OS) compared to high-risk patients in the TCGA training (*P* = 2.229e-04, Fig. 2A) and validation cohorts (*P* = 1.638e-02, Fig. 2B).

**Fig. 2.**
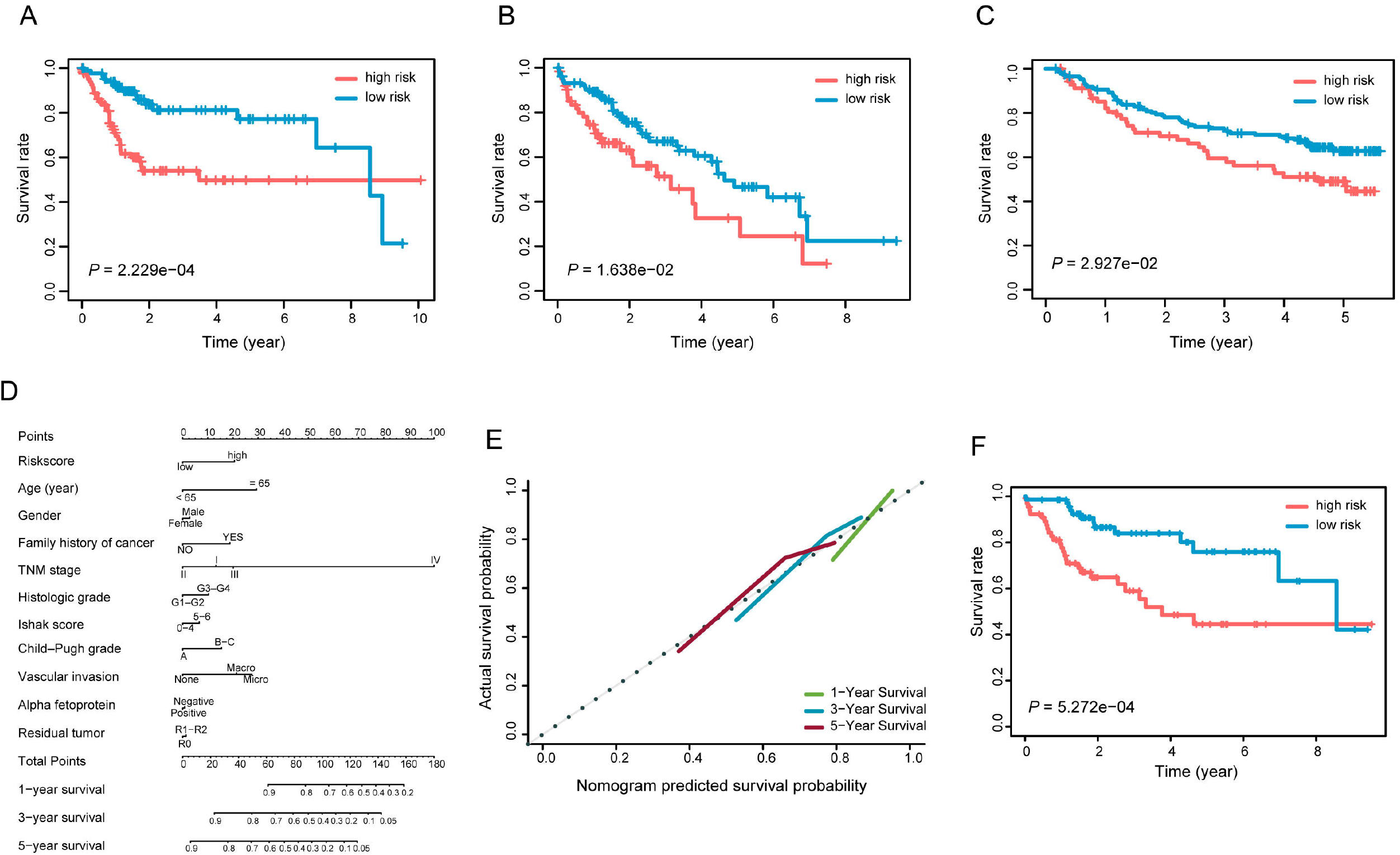
Performance of the predictive gene signature. **A-C**: Kaplan-Meier survival curves for comparison of the overall survival rates between patients in the low-risk group and the high-risk group for the TCGA training cohort (**A**), the TCGA validation cohort (**B**) and the external validation cohort from an independent GEO dataset (**C**). **D**: The nomogram predicting the 1-year, 3-year and 5-year overall survival probability of the HCC patients was created by integrating the gene signature with common clinical features. **E**: The calibration plot showing the predicted and the actual observed survival rates for HCC patients at the 1-year, 3-year and 5-year time points. **F**: Survival curves of the nomogram in predicting overall survival probabilities for HCC patients.

Time-dependent receiver operating characteristic (ROC) curves were applied to evaluate the predicting capacity of the two-E2F prognostic signature. Area under the curve (AUC) yielded by the prognostic signature at 1-year, 3-year and 5-year survival prediction was 0.749, 0.683, and 0.698, respectively, for the TCGA training cohort (Fig. S1A). We further confirmed the prediction efficiency for the two-E2F signature in the validation cohort, and the AUC for 1-year, 3-year and 5-year OS rate was 0.661, 0.641 and 0.659, respectively (Fig. S1B). In addition, it was reassuring to note that the E2F gene signature was able to robustly stratify patients into high- and low-risk groups for OS rates in an external validation cohort (GSE14520, *n* = 222, Fig. 2C). These findings demonstrated the effectiveness of the E2F signature in predicting unfavorable prognosis for HCC patients.

### The prognostic risk score showed independence of conventional clinicopathological parameters

To further evaluate the potential of the E2F prognostic signature, we investigated the relationship between the risk level and the potential clinical outcomes by using the whole set of 339 TCGA patients. We found that a significant difference exists in several clinicopathological factors, such as TNM stage, histologic grade, Ishak score, Child-Pugh grade, vascular invasion, and the serum alpha fetoprotein between the two risk groups. However, the risk score was not significantly associated with age, gender, and family history of cancer (Table 1). We next performed a Cox regression analysis on the E2F signature-derived risk score incorporating the commonly used clinicopathologic parameters. Multivariate Cox analysis included only the features that were statistically significant in the univariate analysis of HCC patients (Table 2).

**Table 1.**
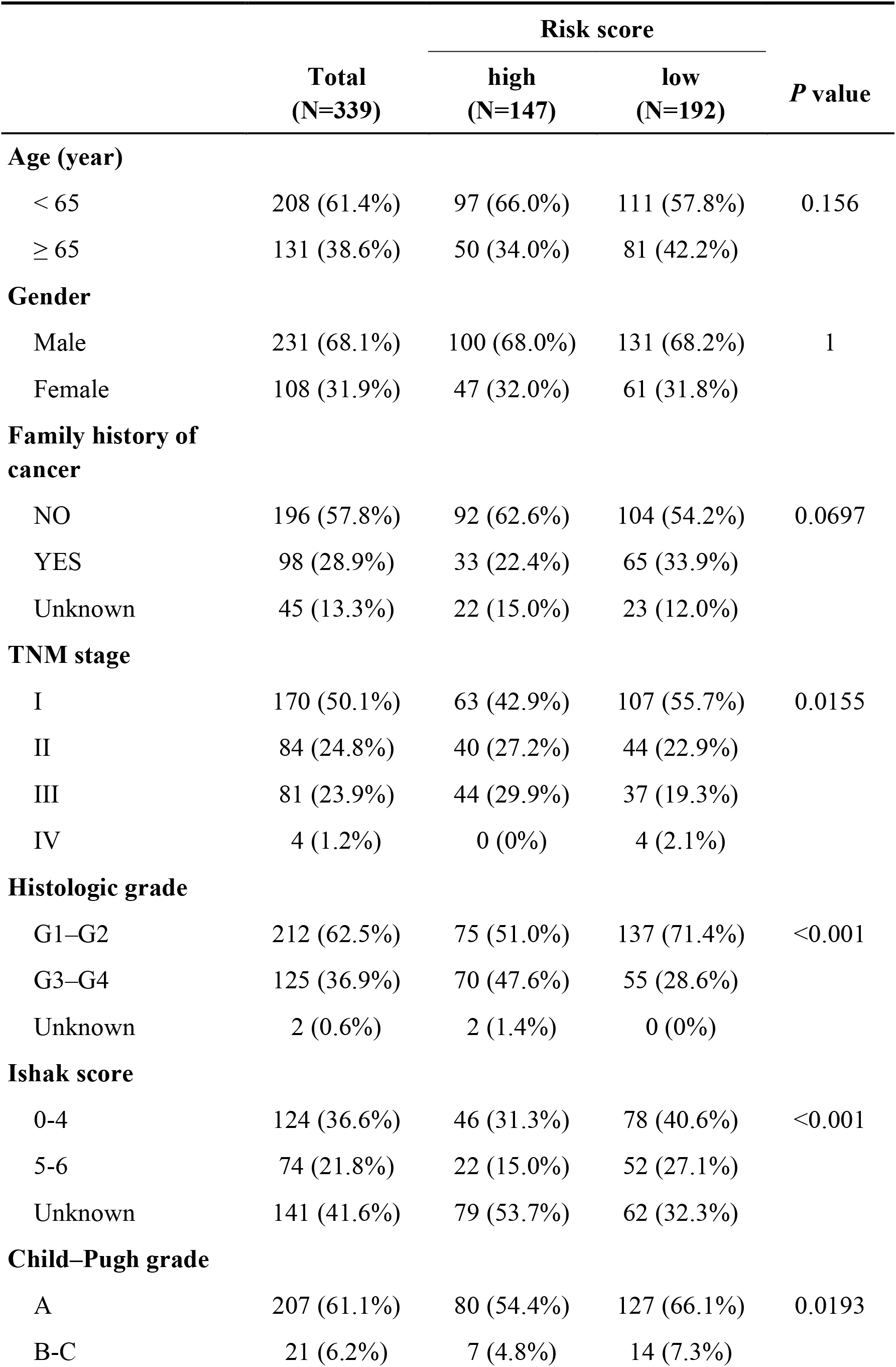
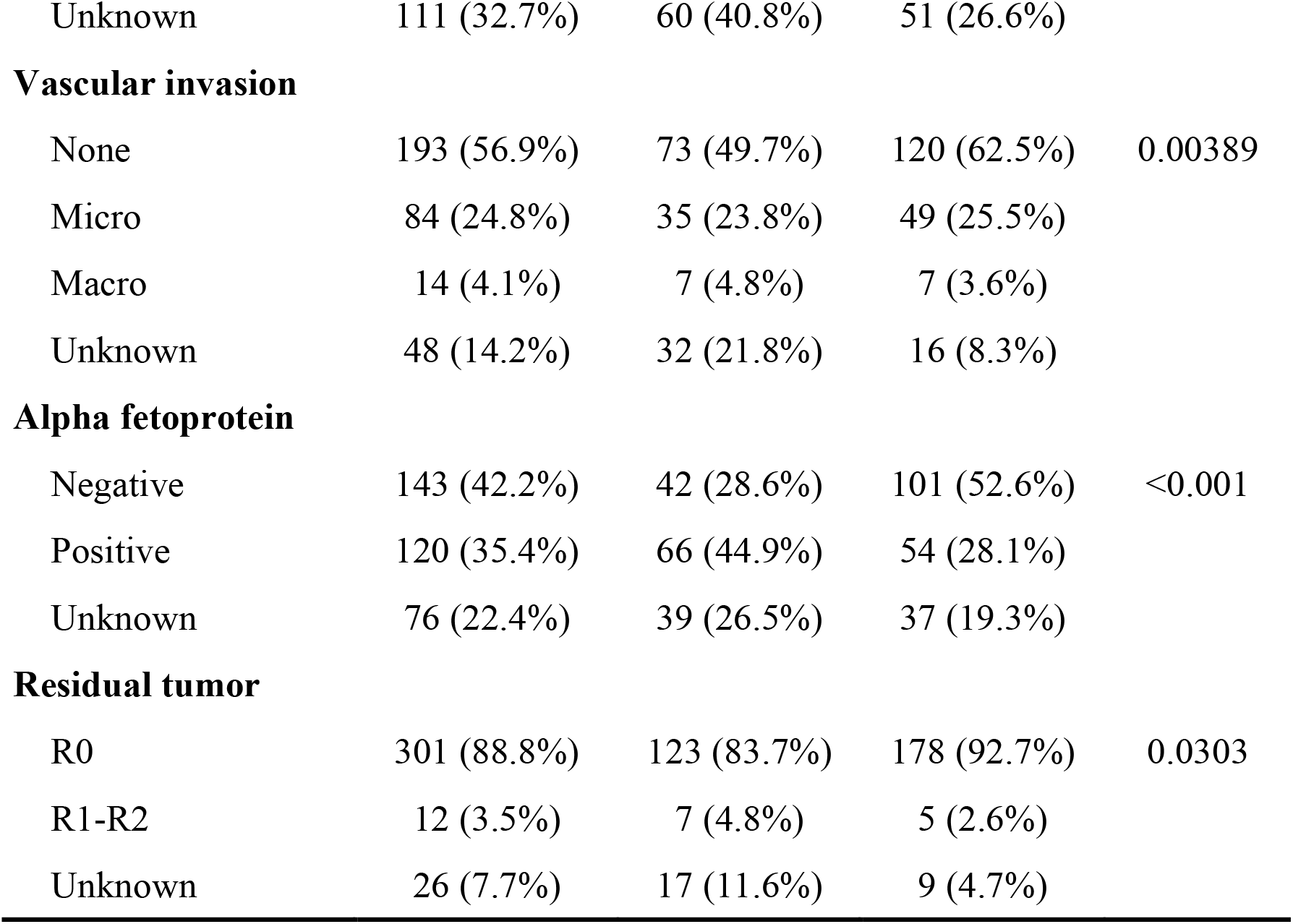
Association between the risk score derived from the prognostic signature and the common clinicopathological factors.

**Table 2.** Univariate and multivariate analysis of overall survival.

| Variables | Univariate analysis |  | Multivariate analysis |  |
| --- | --- | --- | --- | --- |
|  | HR (95% CI) | *P* value | HR (95% CI) | *P* value |
| Age ($\geq 65$ vs. $< 65$) | 1.23(0.85,1.78) | 0.273 | - | - |
| Gender (Female vs. Male) | 1.26(0.87,1.84) | 0.228 | - | - |
| Family history of cancer (YES vs. NO) | 1.14(0.76,1.69) | 0.530 | - | - |
| TNM stage (II vs. I) | 1.42(0.87,2.32) | 0.160 | 1.23(0.66,2.3) | 0.507 |
| TNM stage (III vs. I) | 2.72(1.78,4.15) | **<0.001** | 2.11(1.24,3.59) | **0.006** |
| TNM stage (IV vs. I) | 5.44(1.68,17.63) | **0.005** | 7.3(2.19,24.35) | **0.001** |
| Histologic grade (G3–G4 vs. G1–G2) | 1.14(0.78,1.67) | 0.489 | - | - |
| Ishak score (5–6 vs. 0–4) | 0.87(0.5,1.5) | 0.612 | - | - |
| Child–Pugh grade (B–C vs. A) | 1.66(0.82,3.36) | 0.159 | - | - |
| Vascular invasion (Micro vs. none) | 1.16(0.72,1.88) | 0.539 | 0.97(0.56,1.69) | 0.914 |
| Vascular invasion (Macro vs. none) | 2.52(1.14,5.58) | **0.023** | 1.87(0.82,4.27) | 0.135 |
| Alpha fetoprotein (positive vs. negative) | 1.45(0.92,2.28) | 0.108 | - | - |
| Residual tumor (R1–R2 vs. R0) | 1.17(0.43,3.2) | 0.754 | - | - |
| Risk score (high vs. low) | 2.17(1.5,3.15) | **<0.001** | 1.86(1.19,2.9) | **0.007** |

To provide a clinically adaptable approach to quantifying the risk assessment for HCC patients, we developed a nomogram integrated with the E2F signature and the commonly used clinicopathologic risk factors in the entire TCGA samples dataset (Fig. 2D). We calculated the C-statistic discriminatory index (C-index) and the calibration plot to assess the prognostic performance of the nomogram. As shown in the calibration plot, there was excellent concordance between the predicted outcomes and the observed 1-year, 3-year and 5-year OS (Fig. 2E). Moreover, the C-index for the nomogram was 0.712 (95% CI: 0.683 - 0.742). In addition, a Kaplan-Meier analysis based on the median risk score derived from the nomogram integrating the two-E2F signature and the clinicopathologic features was carried out to measure the discriminatory power of this nomogram. The patients being labeled as high-risk exhibited significantly poorer prognosis than the low-risk patients (Fig. 2F). These findings demonstrated the validity of our E2F signature in combination with the commonly used clinicopathologic factors in predicting survival outcomes for HCC patients.

### Estimation of immune infiltration levels for patients in different risk groups from bulk sequencing data

To elucidate the differences in immune microenvironments between high- and low-risk groups, we applied the CIBERSORT algorithm to infer the fraction of different types of immune cells using the TCGA cohort expression data. The immune cell distribution was calculated based on a matrix of 22 immune cell type signatures (LM22) and compared between high- and low-risk HCC patients (Fig. 3A). The proportion of tumorinfiltrating macrophages M0 and Tregs were significantly higher in the high-risk HCC patients than those in the low-risk group, while the low-risk patients had markedly higher proportions of resting memory CD4^+^ T cells than the high-risk HCC patients (Fig. 3B). These results highlighted a potential mechanistic link between heterogeneous immune infiltration status across different HCC patients and variable clinical manifestations.

**Fig. 3.**
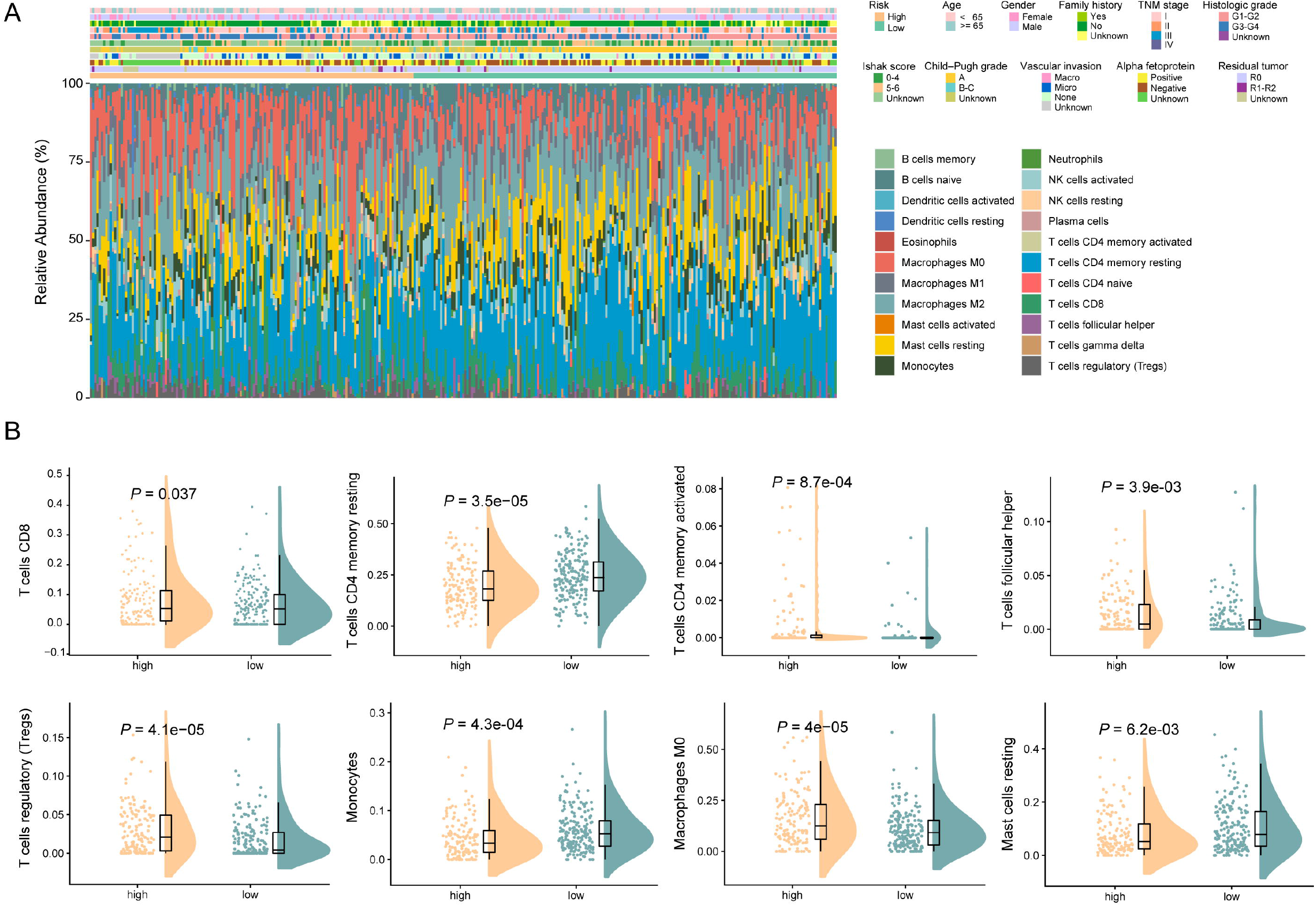
Deconvolution of bulk sequencing data showed the immune cell infiltration landscape in HCC patients from the high-risk and low-risk groups. **A**: Distribution in fractions of diverse tumor infiltrating immune cells in high-risk and low-risk HCC patients. **B**: Comparisons of the proportions of different tumor infiltrating immune cells between the high-risk and low-risk patients. Only the comparisons with statistical significance were visualized.

**Fig. 4.**
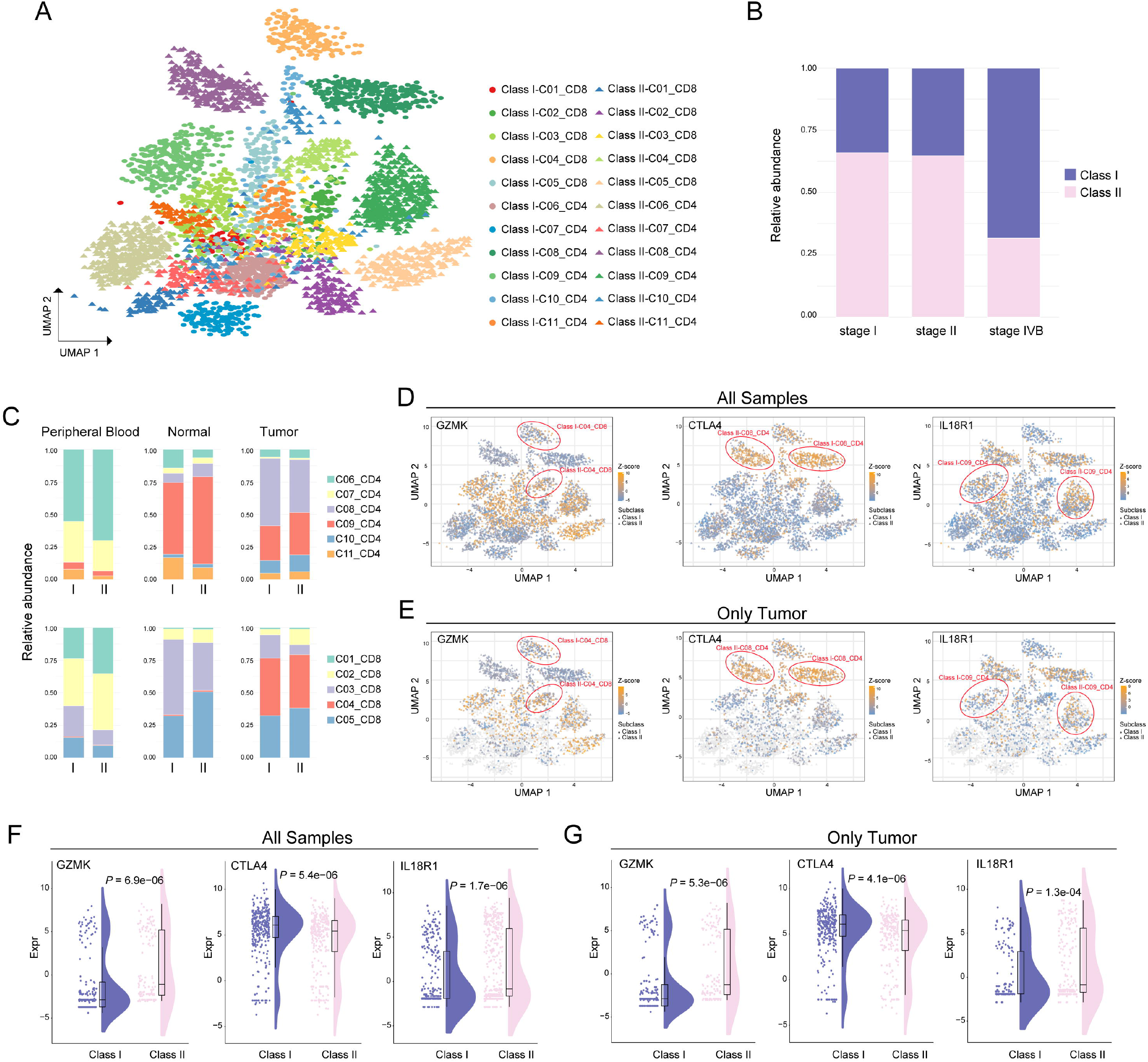
Discriminating T cell populations based on single cell RNA sequencing profiles. **A**: Uniform Manifold Approximation and Projection (UMAP) plot displaying 22 T cell subpopulations identified by the predictive gene model based on the existing 11 clusters. Each dot represents one single cell. The existing 11 clusters: C01_CD8, naive CD8^+^ T cells; C02_CD8, effector CD8^+^ T cells; C03_CD8, mucosal-associated invariant T cells (MAIT); C04_CD8, exhausted CD8^+^ T cells; C05_CD8, intermediate state CD8^+^ T cells between effector and exhausted CD8^+^ T cells; C06_CD4, naive CD4^+^ T cells; C07_CD4, peripheral T regulatory cells (Tregs); C08_CD4, tumor Tregs; C09_CD4, mixed state cells; C10_CD4, exhausted CD4^+^ T cells; C11_CD4, cytotoxic CD4^+^ T cells. **B**: The proportions of Class I and Class II subpopulations in each stage of the HCC patients. **C**: The proportions of each T cell type between the Class I and Class II populations across peripheral blood, tumor and adjacent normal tissues. **D**: UMAP plot of all T cells from peripheral blood, normal and tumor tissues, with all cells colored according to the gene expression of GZMK, CTLA4 and IL18R1, respectively. The two cell subpopulations between which the respective gene was differentially expressed were marked with red circles. **E**: UMAP plot of all T cells, with only tumor cells colored according the gene expression of GZMK, CTLA4 and IL18R1, respectively. Non-tumor cells were colored gray. The two cell subpopulations between which the respective gene was differentially expressed were marked with red circles. **F**: Differential expression of GZMK, CTLA4 and IL18R1 in all cell samples. **G**: Differential expression of GZMK, CTLA4 and IL18R1 in tumor cell samples.

### Efficacy of the predictive signature in discriminating cell types using single-cell transcriptomic profiles

Deconvolution of bulk transcriptomes implemented with CIBERSORT showed remarkable intertumoral immune cell heterogeneity between patients from different risk groups. It is possible that the gene signature developed initially to stratify HCC patients with different outcome risk could discriminate infiltrating T cells in the tumor microenvironment. To explore this hypothesis, we extended the gene signature on a published single-cell RNA sequencing dataset with 4,070 T cell samples of six HCC patients (5), with the classification being applied to single-cell objects instead of traditional patient samples (Fig. 4A). We subcategorized the infiltrating T cells into two classes (Class I and II) based on the E2F signature (Figs. 4A and 4B) (5). Surprisingly, we observed that late-stage HCC patients exhibited a significantly higher abundance of Class I cells than patients from other stages (Fig. 4B, *P* = 2.12e-100, Chi-square test).

The composition of immune cells varied spatially between Class I and II for both CD4^+^ and CD8^+^ T cells (Fig. 4C). Cells from the first CD8^+^ T cell cluster (C01_CD8) were characterized as naive CD8^+^ T cells. Cells from the first CD4^+^ T cell cluster (C06_CD4) were characterized as naive CD4^+^ T cells. These were both more abundant in peripheral blood samples than samples from tumor and normal tissues. The class II cells harbored higher abundances of naive CD4^+^ and CD8^+^ T cells than the Class I cells. In tumor tissues, the effector CD8^+^ T cells, C02_CD8, were more prevalent in the Class II than Class I group, while the tumor Tregs (C08_CD4) were higher in Class I subpopulations. For both Class I and II, the exhausted T cells (C04_CD8 and C10_CD4) were more abundant in tumor tissues than peripheral blood and adjacent normal tissues. In addition, the fraction of mucosal-associated invariant T (MAIT) cells (C03_CD8) was higher in Class I than Class II among all peripheral blood, normal, and tumor cell samples. Within the exhausted CD8+ T cells, we observed the expression levels of cytotoxic effector GZMK was markedly lower in Class I subpopulations than Class II subpopulations not only for tumor cell samples but also for T cells from all three locations (Figs. 4D-G). As a well-established exhaustion marker and immune checkpoint molecule, CTLA4 exhibited significantly higher expression levels in Class I tumor Tregs than Class II tumor Tregs (Figs. 4D-G). Interestingly, among cells in the C09_CD4 cluster, which were cells primarily in an intermediate state between cytolytic and suppressive T cells, we observed significantly lower expression levels of IL18R1 in Class I subpopulations than Class II group (Figs. 4D-G), which might be associated with the process of transition to exhausted T cells (22, 23).

### Distinct cell fates revealed between Class I and Class II subpopulations

To further explore the developmental fates of Class I and II cells, next we performed pseudotime analyses using Monocle (20) to infer the dynamic cell states and immune cell transitions for both subclasses in CD4^+^ and CD8^+^ T cells. For CD8^+^ T cells, we excluded the MAIT cells because of their distinct characteristics of TCRs (5). Monocle pseudotime analysis showed that Class I and II subpopulations exhibited different developmental trajectories and cell type distributions for both CD4^+^ and CD8^+^ T cells (Fig. 5A and Fig. S2). We observed that the trajectory path of Class I subpopulations in CD4^+^ cells contained only one branch point while the trajectory path of Class II subpopulations from naive to exhausted CD4^+^ T cells contained two branch points. The Class I subpopulations of naive and cytotoxic CD4^+^ T cells (Class I-C06_CD4 and Class I-C11_CD4) overlapped extensively and resided mostly at the beginning state of the pseudotime path (Fig. 5A, upper left), whereas the developmental fates of these two cell types differed markedly in Class II CD4^+^ subpopulations, with the cytotoxic subpopulations (Class II-C11_CD4) localized mainly at a distal branch (Fig. 5A, lower left). For CD8^+^ T cells, the Class II subpopulations with an intermediate status between effector and exhausted phenotypes (Class II-C05_CD8) distributed uniformly along pseudotime and overlapped with the exhausted CD8^+^ T cells (Class II-C04_CD8) along the trajectory path, especially at the terminal state (Fig. 5A, lower right). Unlike the Class II subpopulations, when the trajectory bifurcated into two branches in Class I subpopulations, the Class I-C05_CD8 cells within the intermediate state almost disappeared and the two branches generated after the developmental branch point were enriched for exhausted CD8^+^ T cells (Class I-C04_CD8), which emerged earlier than those in Class II subpopulations (Fig. 5A, upper right).

**Fig. 5.**
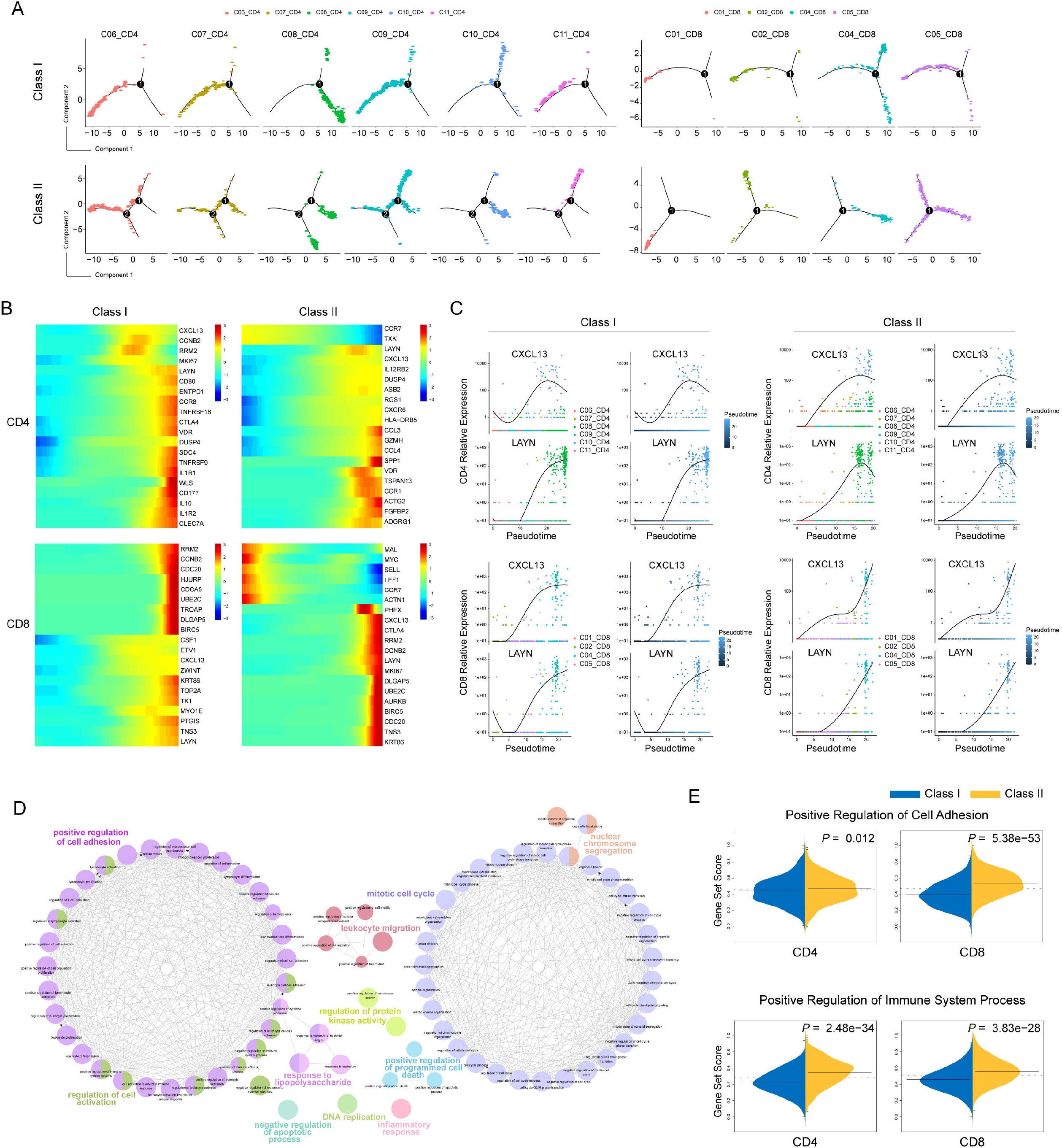
Different developmental cell fates between the Class I and Class II cells. **A**: Faceted pseudotime plot according to T cell types for the Class I and Class II subpopulations. **B**: Heatmap plot for the expression of the 20 most dynamic genes along pseudotime for Class I and Class II subpopulations. **C**: The dynamic expression patterns of the marker genes CXCL13 and LAYN in Class I and Class II subpopulations. **D**: Gene Ontology (GO) results showed the biological processes that associated with the significantly dynamic genes from the branch-dependent expression analysis. Each node represents an enriched GO term. **E**: ssGSEA results for comparing the expression of the dynamic genes during branch evolution for the “Positive Regulation of Cell Adhesion” term and the “Positive Regulation of Immune System Process” term.

We then explored the transcriptional changes along the pseudotime trajectory and identified genes that were significantly dynamic during the differentiation processes (Fig. 5B). The osteopontin protein encoded by the SPP1 gene, ranked as the most significantly associated factor with the Class II CD4^+^ T cells and upregulated prominently at the terminal state (Tables S4-S7, Fig. S3), has been demonstrated to be a potential driver of tumor cell biodiversity (6). Among each set of the top 20 genes that were significantly associated with the transitional states along pseudotime, the T cell recruitment chemokine CXCL13 and layilin (LAYN) were common genes for all four subpopulations of CD4^+^ and CD8^+^ T cells (Figs. 5B and 5C). CXCL13 serves as an exhaustion marker and is also a crucial factor involved in the formation of tertiary lymphoid structures. The secretion of CXCL13 has been shown to participate in the exhaustion of CD4^+^ T cells and the progression of CD8^+^ T cells towards late dysfunctionality (24). CXCL13 exhibited similar expression patterns in Class I and II CD4+ T cells, whereas the Treg and exhaustion marker LAYN showed a differential trend, with a modest upregulation observed at the terminal end of the trajectory in Class I CD4+ T cells compared to the noticeable reduction in the Class II CD4+ subpopulation (Fig. 5C). We also observed that within the Class II subpopulations of the CD8^+^ T cells, the expression of CXCL13 increased dramatically at the end stage of the trajectory, corresponding to the exhausted cell state after a relatively steady phase of the intermediate cell state. Unlike CXCL13 which has a stationary point, the consistently elevated expression of LAYN was observed for the whole differentiation processes of the Class II CD8^+^ T cells (Fig. 5C). As a suppressive marker, LAYN also showed a powerful capacity for discriminating HCC patients with different survival outcomes (5). Interestingly, we found substantial overlap between top-ranked genes of Class I and II CD8^+^ T cells, such as cell cycle-regulated molecules RRM2, CCNB2, UBE2C, DLGAP5, CDC20, and BIRC5, as well as the cell adhesion protein TNS3 (Fig. 5B, Tables S4-S7). The cell cycle genes showed similar patterns of expression in Class I and II CD8^+^ T cells, whereas TNS3, CXCL13, and LAYN exhibited inconsistent trend during the differentiation processes (Fig. 5B), suggesting that the adhesion and migration levels could differ between the Class I and II CD8^+^ T cells (25–27).

To further delineate the processes represented by the developmental trajectory, we then carried out a branch expression analysis to identify the branch-dependent transcriptional and functional differences between the two subpopulations of CD4^+^ and CD8^+^ T cells (Fig. S4, Tables S8-S12). We explored the genes that were significantly associated with each cell fate decision point for the subpopulations in the CD4^+^ and CD8^+^ T cells and found that the top 100 genes responsible for the cell fate decisions were overrepresented by several important biological processes, including the positive regulation of cell adhesion and mitotic cell cycle (Fig. 5D, Table S13). To investigate whether a similar profile of genes exerted different effects in developmental fate decisions among different cell subpopulations, we carried out a single sample gene set enrichment analysis (ssGSEA) to compare transcriptional activities between Class I and II subpopulations. We observed that the gene set scores were remarkably higher in the Class II-CD8 subpopulation than the Class I-CD8 subpopulation for the “Positive Regulation of Cell Adhesion” and “Positive Regulation of Immune System Process” categories (Fig. 5E), suggesting that the Class II-CD8 cells may represent a T cell subset with a higher cytotoxicity than the Class I-CD8 cells. In addition, the cell cycle genes showed a significantly higher enrichment score in Class I subpopulations for both CD4^+^ and CD8^+^ T cells (Fig. S5), supporting the role of tumor microenvironmental reprogramming in cell cycle perturbation (28).

## Discussion

During the progression of the cell cycle, the regulation of cell proliferation involves a cascade of molecular interactions associated with genome stability, DNA repair, and chromatin regulation (29). Mechanistic studies have exposed a pivotal role for the CDK-RB-E2F axis in the maintenance of genomic integrity and cell homeostasis, as well as in responding to DNA damage and stress (30–33). The CDK inhibitors have long been regarded as promising targets for therapeutic intervention in cancer, mediating the phosphorylation and activation of RB proteins and thereby enabling the release of E2F transcription factors that regulate a plethora of genes required for DNA synthesis and chromosome stability during cell cycle progression (34–37). An unequivocal consensus has been reached on the central role of the CDK-RB-E2F pathway in cancer (38, 39). The E2Fs represent a crucial group of transcriptional regulators that are involved in cell cycle control and consists of nine different gene products encoded by eight chromosomal loci. Based on *in vitro* structural studies, the E2F family members have been artificially subdivided into three categories according to the sequence homology and protein function (40, 41), although challenged by *in vivo* studies as being a gross oversimplification (42, 43). While the precise mechanisms remain unclear and warrant further research, dysregulated functions of the E2F family members and/or the target genes associated with apoptosis, metabolism, or angiogenesis have been linked to poor outcomes in cancers (12, 16–18). In this study, we established a prognostic signature based on the expression profiles of the E2F transcription factors that could stratify patients with HCC into different risk groups. Patients classified into the high-risk group exhibited significantly more unfavorable prognosis than the low-risk patients. As the component genes in the signature, E2F2 and E2F5 may act in combination to coordinate a series of cellular processes involved in cancer aggressiveness.

E2F2 is traditionally classified as an activator and it has been reported that the E2F2 gene plays a pivotal role in liver regeneration, the absence of which results in the deregulation of S-phase entry and inhibition of hepatocytes proliferation in mice (44). Previous studies have demonstrated the contributory role of E2F2 upregulation in the development of breast cancer, owing to the enrichment of its target genes related to angiogenesis and extracellular matrix remodeling (45). Furthermore, E2F2 also possesses pro-apoptotic functions during DNA damage (46). In HCC, there has been evidence suggesting that the down-regulation of E2F2 could result in the inhibition of hepatocyte proliferation and prevention of cell progression (47). Rather than being a traditionally acknowledged activator as E2F2, E2F5 has been grouped into the repressor group based on previous *in vitro* studies (42, 48). It has been demonstrated that the proliferation and migration of cells in breast cancer can be inhibited by decreasing the transcriptional output of E2F5 through a microRNA-dependent mechanism (49). In addition, E2F5 was also found to be substantially overexpressed in glioblastoma and prostate cancer (50, 51). Furthermore, several studies have sought to explore the relationship between E2F5 and HCC, and showed that the dysregulation of E2F5 expression was associated with the process of tumor development (52, 53).

During the process of tumor cell proliferation and tumor growth, a disturbed immune microenvironment could be involved in promoting tumoral immune escape (54). Immune microenvironment heterogeneity can lead to unresponsiveness and treatment resistance in conventional chemotherapy and targeted immunotherapy (55). Therefore, understanding the cellular and molecular heterogeneity of tumor infiltrating cells from a different perspective may be of crucial importance for making improvements in diagnosis and treatment. Developing gene signatures have been applied widely and evolved rapidly for predicting the prognosis of patients. Our study built and validated a prognostic gene signature based on E2F transcription factors owing to their essential role in HCC as well as in the cell cycle (14, 21). We noted that Huang et al. (56) performed multivariate analysis integrating E2Fs and several clinical parameters and found E2F5 and E2F6 were independent predictive factors corroborating the role of E2Fs in HCC etiology. Notably, the results from Huang et al. (56) does not contradict ours because we built the gene signature merely based on the E2Fs without clinical parameters. Besides, the performance of the combination of E2F5 and E2F6 in predicting the clinical outcomes of HCC patients was not shown.

Our study focused on the delineation of cellular and molecular heterogeneity from the perspective of a prognostic gene signature and attempted to unveil key determinants underlying the capability of the predictive gene signature, although perhaps not the one with the best performance, which requires further study to be identified. The single cells harbored by patients with high expression of the gene signature did not necessarily all show high gene signature expression. However, the presence of enough single cells with high signature expression caused the corresponding patient to be identified as having a high expression of the gene signature and classified as having a high risk of poor survival. The results of our study linked patient prognosis to intratumoral cellular and molecular heterogeneity, highlighted the distinct cell populations in patients across different disease stages, and revealed specific subpopulations of CD8^+^ T cells exhibiting differential cell adhesion level which might result in different tumor dissemination patterns. Investigations on inter-patient heterogeneity and a more comprehensive characterization of cell populations, especially those incorporating transcriptome profiles, clinical information and single-cell data from the same group of individuals will be conducted in future studies.

## DATA AVAILABILITY STATEMENT

Publicly available datasets were analyzed in this study. The data can be found here: TCGA https://www.cancer.gov/about-nci/organization/ccg/research/structural-genomics/tcga and GEO https://www.ncbi.nlm.nih.gov/geo/ repositories (GSE76427, GSE136247, GSE107170, GSE102079, and GSE14520 for bulk-seq data; GSE98638 for single-cell sequencing data).

## AUTHOR CONTRIBUTIONS

X. L., Z. S. and F. M. conceived and designed the study. L. W., F. M. and X. L. wrote the manuscript. L. W. analyzed the data. Y C. revised the manuscript.

## REFERENCES

1. European Association for the Study of the Liver. Electronic address eee, European Association for the Study of the L. EASL Clinical Practice Guidelines: Management of hepatocellular carcinoma. Journal of hepatology (2018) 69(1):182–236. doi: 10.1016/j.jhep.2018.03.019.

2. Ganne-Carrie N, Nahon P. Hepatocellular carcinoma in the setting of alcohol-related liver disease. Journal of hepatology (2019) 70(2):284–93. doi: 10.1016/j.jhep.2018.10.008.

3. Finn RS, Ryoo BY, Merle P, Kudo M, Bouattour M, Lim HY, et al. Pembrolizumab As Second-Line Therapy in Patients With Advanced Hepatocellular Carcinoma in KEYNOTE-240: A Randomized, Double-Blind, Phase III Trial. J Clin Oncol (2020) 38(3):193–202. doi: 10.1200/JCO.19.01307.

4. Sangro B, Gomez-Martin C, de la Mata M, Inarrairaegui M, Garralda E, Barrera P, et al. A clinical trial of CTLA-4 blockade with tremelimumab in patients with hepatocellular carcinoma and chronic hepatitis C. Journal of hepatology (2013) 59(1):81–8. doi: 10.1016/j.jhep.2013.02.022.

5. Zheng C, Zheng L, Yoo JK, Guo H, Zhang Y, Guo X, et al. Landscape of Infiltrating T Cells in Liver Cancer Revealed by Single-Cell Sequencing. Cell (2017) 169(7):1342–56 e16. doi: 10.1016/j.cell.2017.05.035.

6. Ma L, Wang L, Khatib SA, Chang CW, Heinrich S, Dominguez DA, et al. Single-cell atlas of tumor cell evolution in response to therapy in hepatocellular carcinoma and intrahepatic cholangiocarcinoma. Journal of hepatology (2021) 75(6):1397–408. doi: 10.1016/j.jhep.2021.06.028.

7. Zhang Q, He Y, Luo N, Patel SJ, Han Y, Gao R, et al. Landscape and Dynamics of Single Immune Cells in Hepatocellular Carcinoma. Cell (2019) 179(4):829–45 e20. doi: 10.1016/j.cell.2019.10.003.

8. Malato Y, Naqvi S, Schurmann N, Ng R, Wang B, Zape J, et al. Fate tracing of mature hepatocytes in mouse liver homeostasis and regeneration. J Clin Invest (2011) 121(12):4850–60. doi: 10.1172/JCI59261.

9. Hassan M, Ghozlan H, Abdel-Kader O. Activation of RB/E2F signaling pathway is required for the modulation of hepatitis C virus core protein-induced cell growth in liver and non-liver cells. Cell Signal (2004) 16(12):1375–85. doi: 10.1016/j.cellsig.2004.04.005.

10. Mayhew CN, Carter SL, Fox SR, Sexton CR, Reed CA, Srinivasan SV, et al. RB loss abrogates cell cycle control and genome integrity to promote liver tumorigenesis. Gastroenterology (2007) 133(3):976–84. doi: 10.1053/j.gastro.2007.06.025.

11. Kent LN, Leone G. The broken cycle: E2F dysfunction in cancer. Nat Rev Cancer (2019) 19(6):326–38. doi: 10.1038/s41568-019-0143-7.

12. Chen HZ, Tsai SY, Leone G. Emerging roles of E2Fs in cancer: an exit from cell cycle control. Nat Rev Cancer (2009) 9(11):785–97. doi: 10.1038/nrc2696.

13. Attwooll C, Lazzerini Denchi E, Helin K. The E2F family: specific functions and overlapping interests. EMBO J (2004) 23(24):4709–16. doi: 10.1038/sj.emboj.7600481.

14. Rowland BD, Bernards R. Re-evaluating cell-cycle regulation by E2Fs. Cell (2006) 127(5):871–4. doi: 10.1016/j.cell.2006.11.019.

15. Hernando E, Nahle Z, Juan G, Diaz-Rodriguez E, Alaminos M, Hemann M, et al. Rb inactivation promotes genomic instability by uncoupling cell cycle progression from mitotic control. Nature (2004) 430(7001):797–802. doi: 10.1038/nature02820.

16. Lan W, Bian B, Xia Y, Dou S, Gayet O, Bigonnet M, et al. E2F signature is predictive for the pancreatic adenocarcinoma clinical outcome and sensitivity to E2F inhibitors, but not for the response to cytotoxic-based treatments. Sci Rep (2018) 8(1):8330. doi: 10.1038/s41598-018-26613-z.

17. Kent LN, Rakijas JB, Pandit SK, Westendorp B, Chen HZ, Huntington JT, et al. E2f8 mediates tumor suppression in postnatal liver development. J Clin Invest (2016) 126(8):2955–69. doi: 10.1172/JCI85506.

18. Kent LN, Bae S, Tsai SY, Tang X, Srivastava A, Koivisto C, et al. Dosage-dependent copy number gains in E2f1 and E2f3 drive hepatocellular carcinoma. J Clin Invest (2017) 127(3):830–42. doi: 10.1172/JCI87583.

19. Leek JT, Johnson WE, Parker HS, Jaffe AE, Storey JD. The sva package for removing batch effects and other unwanted variation in high-throughput experiments. Bioinformatics (2012) 28(6):882–3. doi: 10.1093/bioinformatics/bts034.

20. Qiu X, Mao Q, Tang Y, Wang L, Chawla R, Pliner HA, et al. Reversed graph embedding resolves complex single-cell trajectories. Nat Methods (2017) 14(10):979–82. doi: 10.1038/nmeth.4402.

21. Zhan L, Huang C, Meng XM, Song Y, Wu XQ, Miu CG, et al. Promising roles of mammalian E2Fs in hepatocellular carcinoma. Cell Signal (2014) 26(5):1075–81. doi: 10.1016/j.cellsig.2014.01.008.

22. Ingram JT, Yi JS, Zajac AJ. Exhausted CD8 T cells downregulate the IL-18 receptor and become unresponsive to inflammatory cytokines and bacterial co-infections. PLoS Pathog (2011) 7(9):e1002273. doi: 10.1371/journal.ppat.1002273.

23. Okamura H, Tsutsi H, Komatsu T, Yutsudo M, Hakura A, Tanimoto T, et al. Cloning of a new cytokine that induces IFN-gamma production by T cells. Nature (1995) 378(6552):88–91. doi: 10.1038/378088a0.

24. van der Leun AM, Thommen DS, Schumacher TN. CD8(+) T cell states in human cancer: insights from single-cell analysis. Nat Rev Cancer (2020) 20(4):218–32. doi: 10.1038/s41568-019-0235-4.

25. Mahuron KM, Moreau JM, Glasgow JE, Boda DP, Pauli ML, Gouirand V, et al. Layilin augments integrin activation to promote antitumor immunity. J Exp Med (2020) 217(9). doi: 10.1084/jem.20192080.

26. Vos JM, Tsakmaklis N, Patterson CJ, Meid K, Castillo JJ, Brodsky P, et al. CXCL13 levels are elevated in patients with Waldenstrom macroglobulinemia, and are predictive of major response to ibrutinib. Haematologica (2017) 102(11):e452–e5. doi: 10.3324/haematol.2017.172627.

27. Bono P, Rubin K, Higgins JM, Hynes RO. Layilin, a novel integral membrane protein, is a hyaluronan receptor. Mol Biol Cell (2001) 12(4):891–900. doi: 10.1091/mbc.12.4.891.

28. Powathil GG, Gordon KE, Hill LA, Chaplain MA. Modelling the effects of cell-cycle heterogeneity on the response of a solid tumour to chemotherapy: biological insights from a hybrid multiscale cellular automaton model. J Theor Biol (2012) 308:1–19. doi: 10.1016/j.jtbi.2012.05.015.

29. Otto T, Sicinski P. Cell cycle proteins as promising targets in cancer therapy. Nat Rev Cancer (2017) 17(2):93–115. doi: 10.1038/nrc.2016.138.

30. Ohtani N, Brennan P, Gaubatz S, Sanij E, Hertzog P, Wolvetang E, et al. Epstein-Barr virus LMP1 blocks p16INK4a-RB pathway by promoting nuclear export of E2F4/5. J Ce**l**l Biol (2003) 162(2):173–83. doi: 10.1083/jcb.200302085.

31. Wetmore C, Boyett J, Li S, Lin T, Bendel A, Gajjar A, et al. Alisertib is active as single agent in recurrent atypical teratoid rhabdoid tumors in 4 children. Neuro Oncol (2015) 17(6):882–8. doi: 10.1093/neuonc/nov017.

32. Kastan MB, Bartek J. Cell-cycle checkpoints and cancer. Nature (2004) 432(7015):316–23. doi: 10.1038/nature03097.

33. Bartek J, Lukas C, Lukas J. Checking on DNA damage in S phase. Nat Rev Mol Cell Biol (2004) 5(10):792–804. doi: 10.1038/nrm1493.

34. Hutcheson J, Witkiewicz AK, Knudsen ES. The RB tumor suppressor at the intersection of proliferation and immunity: relevance to disease immune evasion and immunotherapy. Cell Cycle (2015) 14(24):3812–9. doi: 10.1080/15384101.2015.1010922.

35. Malumbres M, Barbacid M. Mammalian cyclin-dependent kinases. Trends Biochem Sci (2005) 30(11):630–41. doi: 10.1016/j.tibs.2005.09.005.

36. Sherr CJ, Roberts JM. Living with or without cyclins and cyclin-dependent kinases. Genes Dev (2004) 18(22):2699–711. doi: 10.1101/gad.1256504.

37. Hydbring P, Malumbres M, Sicinski P. Non-canonical functions of cell cycle cyclins and cyclin-dependent kinases. Nat Rev Mol Cell Biol (2016) 17(5):280–92. doi: 10.1038/nrm.2016.27.

38. Sherr CJ. Cancer cell cycles. Science (1996) 274(5293):1672–7. doi: 10.1126/science.274.5293.1672.

39. Sherr CJ, McCormick F. The RB and p53 pathways in cancer. Cancer Cell (2002) 2(2):103–12. doi: 10.1016/s1535-6108(02)00102-2.

40. Lammens T, Li J, Leone G, De Veylder L. Atypical E2Fs: new players in the E2F transcription factor family. Trends Cell Biol (2009) 19(3):111–8. doi: 10.1016/j.tcb.2009.01.002.

41. Iaquinta PJ, Lees JA. Life and death decisions by the E2F transcription factors. Curr Opin Cell Biol (2007) 19(6):649–57. doi: 10.1016/j.ceb.2007.10.006.

42. Chen HZ, Ouseph MM, Li J, Pecot T, Chokshi V, Kent L, et al. Canonical and atypical E2Fs regulate the mammalian endocycle. Nat Cell Biol (2012) 14(11):1192–202. doi: 10.1038/ncb2595.

43. Chen YL, Uen YH, Li CF, Horng KC, Chen LR, Wu WR, et al. The E2F transcription factor 1 transactives stathmin 1 in hepatocellular carcinoma. Ann Surg Oncol (2013) 20(12):4041–54. doi: 10.1245/s10434-012-2519-8.

44. Delgado I, Fresnedo O, Iglesias A, Rueda Y, Syn WK, Zubiaga AM, et al. A role for transcription factor E2F2 in hepatocyte proliferation and timely liver regeneration. Am J Physiol Gastrointest Liver Physiol (2011) 301(1):G20–31. doi: 10.1152/ajpgi.00481.2010.

45. Hollern DP, Honeysett J, Cardiff RD, Andrechek ER. The E2F transcription factors regulate tumor development and metastasis in a mouse model of metastatic breast cancer. Mol Cell Biol (2014) 34(17):3229–43. doi: 10.1128/MCB.00737-14.

46. Chen D, Chen Y, Forrest D, Bremner R. E2f2 induces cone photoreceptor apoptosis independent of E2f1 and E2f3. Cell Death Differ (2013) 20(7):931–40. doi: 10.1038/cdd.2013.24.

47. Dong Y, Zou J, Su S, Huang H, Deng Y, Wang B, et al. MicroRNA-218 and microRNA-520a inhibit cell proliferation by downregulating E2F2 in hepatocellular carcinoma. Mol Med Rep (2015) 12(1):1016–22. doi: 10.3892/mmr.2015.3516.

48. Tsai SY, Opavsky R, Sharma N, Wu L, Naidu S, Nolan E, et al. Mouse development with a single E2F activator. Nature (2008) 454(7208):1137–41. doi: 10.1038/nature07066.

49. Xu H, Fei D, Zong S, Fan Z. MicroRNA-154 inhibits growth and invasion of breast cancer cells through targeting E2F5. Am J Transl Res (2016) 8(6):2620–30.

50. Fang DZ, Wang YP, Liu J, Hui XB, Wang XD, Chen X, et al. MicroRNA-129-3p suppresses tumor growth by targeting E2F5 in glioblastoma. Eur Rev Med Pharmacol Sci (2018) 22(4):1044–50. doi: 10.26355/eurrev_201802_14387.

51. Li SL, Sui Y, Sun J, Jiang TQ, Dong G. Identification of tumor suppressive role of microRNA-132 and its target gene in tumorigenesis of prostate cancer. Int J Mol Med (2018) 41(4):2429–33. doi: 10.3892/ijmm.2018.3421.

52. Jiang Y, Yim SH, Xu HD, Jung SH, Yang SY, Hu HJ, et al. A potential oncogenic role of the commonly observed E2F5 overexpression in hepatocellular carcinoma. World J Gastroenterol (2011) 17(4):470–7. doi: 10.3748/wjg.v17.i4.470.

53. Zou C, Li Y, Cao Y, Zhang J, Jiang J, Sheng Y, et al. Up-regulated MicroRNA-181a induces carcinogenesis in hepatitis B virus-related hepatocellular carcinoma by targeting E2F5. BMC Cancer (2014) 14:97. doi: 10.1186/1471-2407-14-97.

54. Kim R, Emi M, Tanabe K. Cancer immunoediting from immune surveillance to immune escape. Immunology (2007) 121(1):1–14. doi: 10.1111/j.1365-2567.2007.02587.x.

55. Khatib S, Pomyen Y, Dang H, Wang XW. Understanding the Cause and Consequence of Tumor Heterogeneity. Trends Cancer (2020) 6(4):267–71. doi: 10.1016/j.trecan.2020.01.010.

56. Huang YL, Ning G, Chen LB, Lian YF, Gu YR, Wang JL, et al. Promising diagnostic and prognostic value of E2Fs in human hepatocellular carcinoma. Cancer Manag Res (2019) 11:1725–40. doi: 10.2147/CMAR.S182001.

