## Supplementary material for "Risk modelling of single cell populations revealed the heterogeneity of immune infiltration in hepatocellular carcinoma": Supplementary Materials.docx

**Supplementary Figure Legends**

**Fig. S1.** Evaluation of the prognostic gene signature in the training and validation cohort. **A** and **B**: The time-dependent ROC curves depicting the accuracy of the prognostic gene signature in predicting the 1-year, 3-year and 5-year overall survival for the training cohort (**A**) and the validation cohort (**B**).

**Fig. S2.** Different pseudotime trajectory paths between the Class I and Class II subpopulations. **A** and **B**: The pseudotime trajectory path from cells within the Class I subpopulation (**A**) and Class II subpopulation (**B**) of the CD4^+^ T cells. **C** and **D**: The pseudotime trajectory path from cells within the Class I subpopulation (**C**) and Class II subpopulation (**D**) of the CD8^+^ T cells. Cells are colored according to the pseudotime.

**Fig. S3.** Transcriptional changes of the SPP1 gene in the Class II subpopulation of CD4^+^ T cells based on the pseudotime analysis.

**Fig. S4.** Expression heatmap of the top 100 genes that significantly associated with the bifurcation of the cells. **A-E**: Cell fate expression of the top 100 most dynamic genes for the branch evolution of the Class I CD4^+^ T cells (**A**), the Class II CD4^+^ T cells at branch point 2 (**B**), the Class II CD4^+^ T cells at branch point 1 (**C**), the Class I CD8^+^ T cells (**D**) and the Class II CD8^+^ T cells (**E**).

**Fig. S5.** ssGSEA results comparing the expression of the “Mitotic Cell Cycle” gene set between the Class I and Class II cells.

**Supplementary Tables**

**Table S1**. Differential gene expression analysis using TCGA LIHC RNA-seq data.

**Table S2**. Differential gene expression analysis using merged RNA-seq data from four GEO datasets (GSE76427, GSE136247, GSE107170 and GSE102079).

**Table S3**. Two E2F genes with statistical significance after multivariate analysis.

**Table S4**. Pseudotime analysis for the Class I subpopulation of CD4^+^ T cells.

**Table S5**. Pseudotime analysis for the Class II subpopulation of CD4^+^ T cells.

**Table S6**. Pseudotime analysis for the Class I subpopulation of CD8^+^ T cells.

**Table S7**. Pseudotime analysis for the Class II subpopulation of CD8^+^ T cells.

**Table S8**. Branch-dependent gene expression analysis for the Class I subpopulation of CD4^+^ T cells.

**Table S9**. Branch-dependent gene expression analysis for the Class II subpopulation of CD4^+^ T cells at branch point 1.

**Table S10**. Branch-dependent gene expression analysis for the Class II subpopulation of CD4^+^ T cells at branch point 2.

**Table S11**. Branch-dependent gene expression analysis for the Class I subpopulation of CD8^+^ T cells

**Table S12**. Branch-dependent gene expression analysis for the Class II subpopulation of CD8^+^ T cells

**Table S13**. Overrepresentation analysis for the combined gene set of the top 100 cell fate decision genes within each T cell subpopulation.
