## Supplementary figures and images for "Risk modelling of single cell populations revealed the heterogeneity of immune infiltration in hepatocellular carcinoma"

### Figure S1.tif

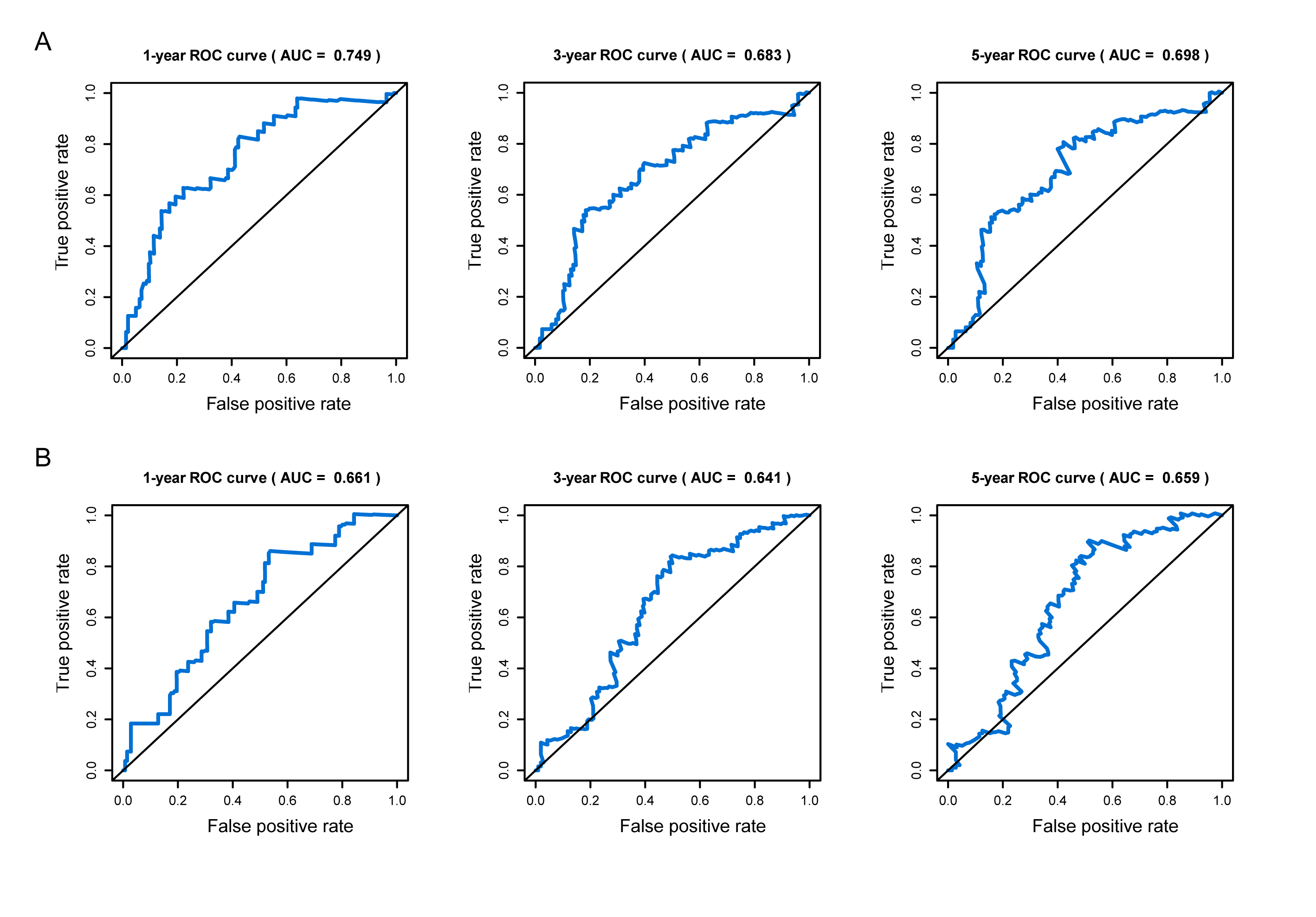

### Figure S2.tif

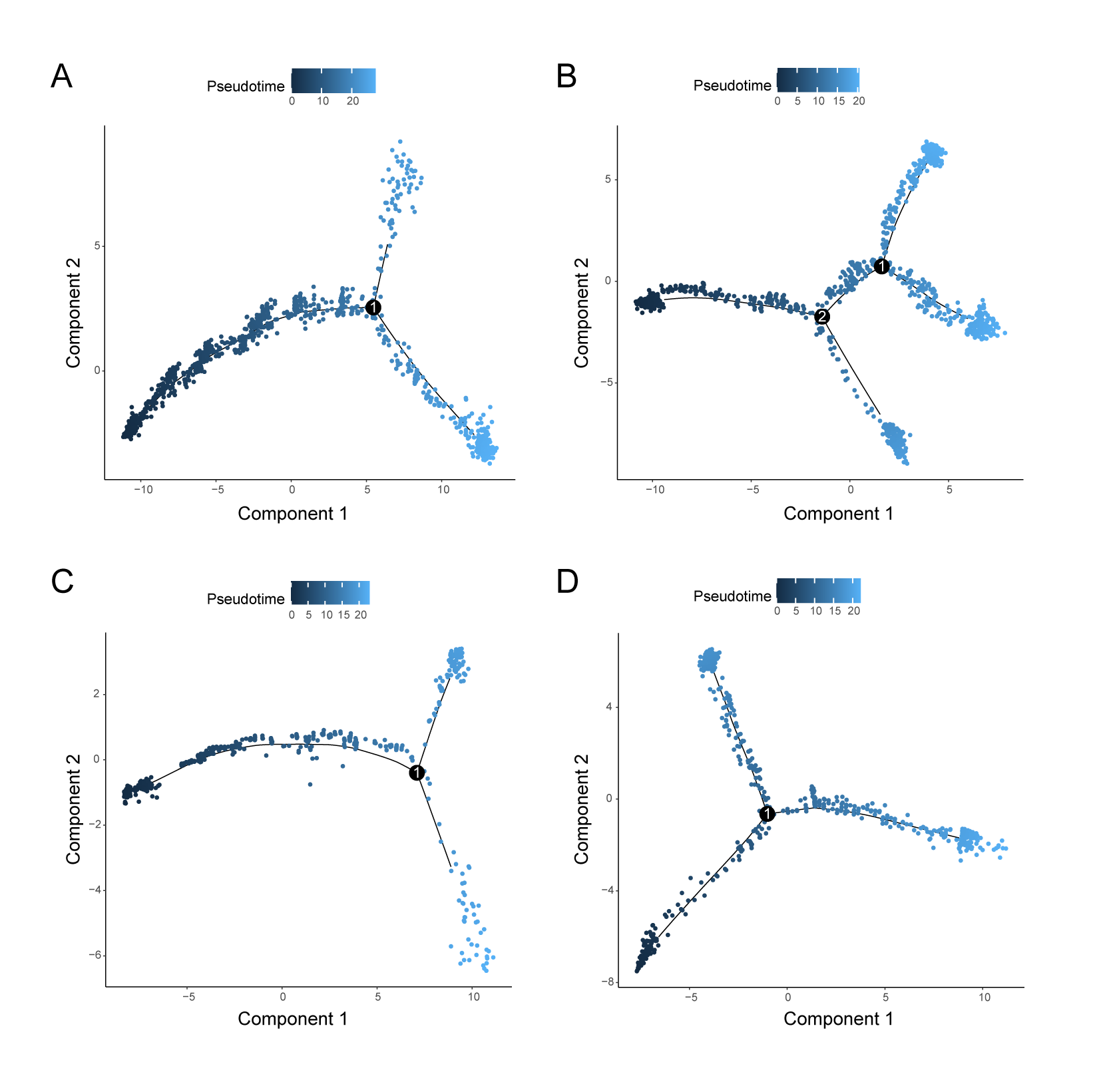

### Figure S3.tif

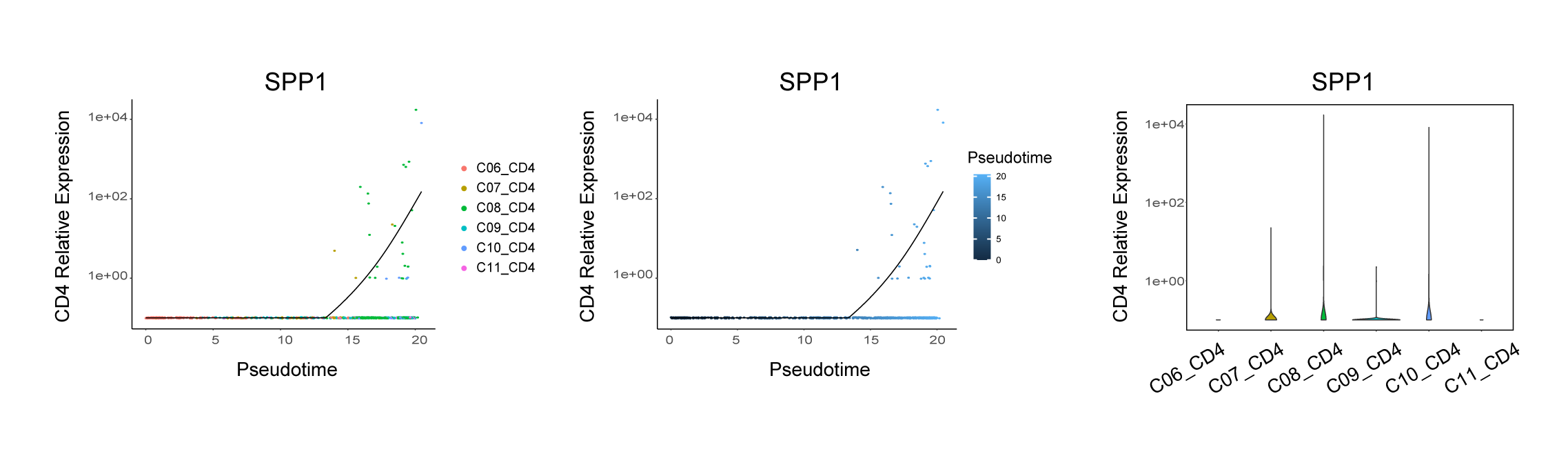

### Figure S4.tif

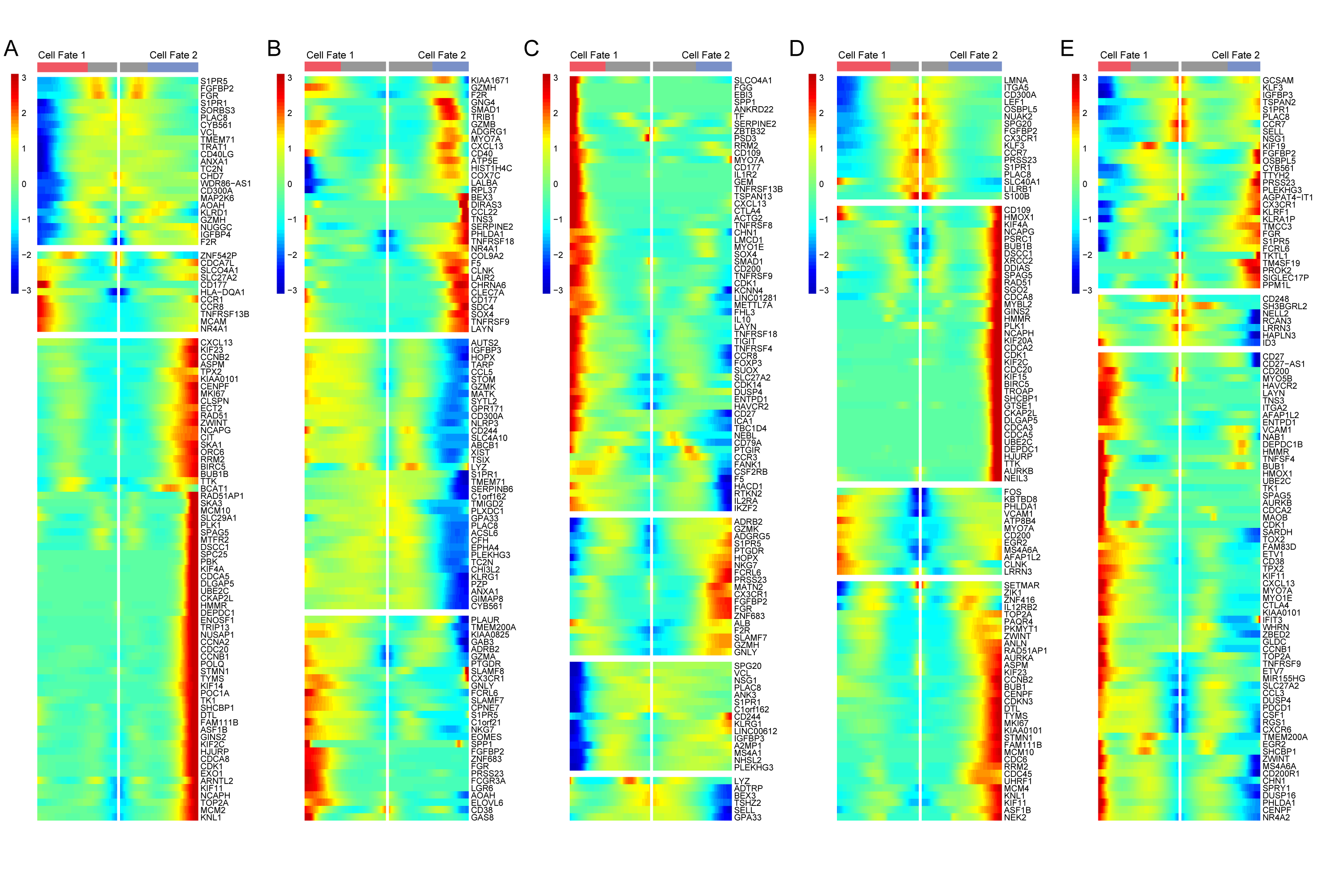

### Figure S5.tif

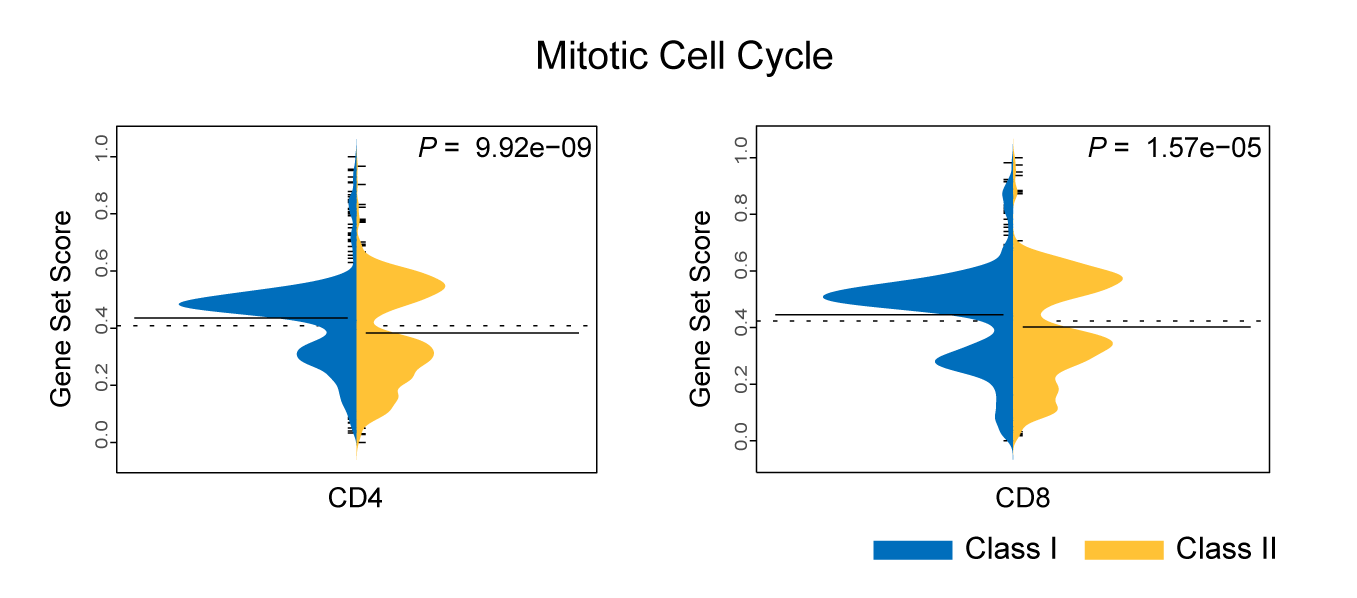
